# Delineating neural contributions to electroencephalogram-based speech decoding

**DOI:** 10.1101/2024.05.09.591996

**Authors:** Motoshige Sato, Yasuo Kabe, Sensho Nobe, Akito Yoshida, Masakazu Inoue, Mayumi Shimizu, Kenichi Tomeoka, Shuntaro Sasai

## Abstract

Speech Brain-computer interfaces (BCIs) have emerged as a pivotal technology in facilitating communication for individuals with speech impairments. Utilizing electroencephalography (EEG) for noninvasive speech BCIs offers an accessible and affordable solution, potentially benefiting a broader audience. However, EEG-based speech decoding remains controversial especially for overt speech, due to difficulties in separating speech-related neural activities from myoelectric potential artifacts generated during articulation. Here we aim to delineate the extent of the neural contributions by employing Explainable AI techniques to a convolutional neural network predicting spoken words based on signals obtained by ultra-high-density (uhd)-EEG. We found that electrode-wise contributions to the decoding cannot be explained by their mutual information with electromyography (EMG). Furthermore, contributing periods of speech to EEG-based decoding are distinct from those to decoding solely relying on EMG. In contrast, there are significant overlaps in signal timings contributing to EEG-based decoding, regardless of vocal conditions such as overt or covert speech. Notably, the denoising process successfully enhanced the decoding contribution from electrodes within speech-related brain areas for all speech conditions. Altogether, our findings support the idea that, with appropriate preprocessing, EEG becomes a valuable tool for decoding spoken words based on underlying neural activities.

## Introduction

Neuroprosthesis technology emerges as a promising solution for individuals with speech impairments due to conditions like brainstem strokes, amyotrophic lateral sclerosis (ALS), or surgeries such as laryngectomy. Traditional communication aids like eye-tracking are significantly slower than natural speech and can lead to fatigue, especially in late-stage ALS where vision and eye movement issues compound these limitations^1–5^. Recent advancements in speech brain-computer interfaces (BCI) through invasive neural recordings, such as intracortical microelectrode arrays or electrocorticogram (ECoG), have shown remarkable potential by achieving word production speeds close to natural conversation^6–11^, presenting a less taxing alternative for users. However, the invasiveness of these methods, which require surgical implantation of electrodes in the brain, poses considerable psychological and physical barriers. This underscores the need for developing speech prostheses with non-invasive neural recording techniques that can overcome the challenge and offer more accessible communication solutions.

Several non-invasive methods have been used in speech decoding, including functional Magnetic Resonance Imaging (fMRI), Magnetoencephalography (MEG), and Electroencephalography (EEG)^12^. fMRI and MEG have greater advantages in their spatial resolutions and have been used to develop successful decoders relevant to speech prostheses^13–15^. However, these devices are large equipment and need shielded rooms in operation, which are unsuitable for practical everyday usage. In contrast, EEG only needs a simpler setting and is more suitable for natural conditions compared to fMRI and MEG. Furthermore, recent innovations in ultra-high-density (uhd)-EEG systems have introduced inter-electrode distances of just 8.6 mm, a significant reduction from the standard 60-65 mm. This advancement may mitigate its spatial resolution limitations and enhance EEG’s viability for speech decoding^16^.

Developing a speech neuroprosthesis needs to collect neural recordings paired with speech. This involves simultaneously recording articulated words and corresponding neural data in overt (articulated) speech paradigms. In contrast, decoding covert (imagined) speech requires different methods to identify produced words, as traditional voices are not used. A common approach is the paced covert production of words, where subjects covertly produce words by accurately following instructed paces. However, the paced production paradigm presents numerous challenges in developing speech prostheses. For example, subjects cannot control the pre-fixed pace, which often differs significantly from their natural word production rates^17,18^. Moreover, this approach obliges subjects to produce pre-determined words or sentences, preventing the capture of subjects’ spontaneous, natural speech. This severely limits the scalability of data collection. Additionally, evidence has shown that the covert speech paradigm substantially reduces decodability^19^, underscoring the need to desterilize the overt speech paradigm in the development of BCIs for covert speech.

To create practical speech BCIs for daily use, extensive efforts have been invested in developing EEG-based decoders. Yet, it remains uncertain whether EEG data collected during overt speech can be repurposed for developing a covert speech decoder. This uncertainty primarily arises because most EEG studies focus on covert speech^12,20–23^ or the moments before articulation onset^24–28^ to avoid contamination from strong EMGs generated by articulation. Consequently, the potential of EEG data during overt speech to reveal neural signatures essential for speech production, and their relevance to covert speech, is not well understood. In this study, we aim to explore the relationship between overt and covert speech. We obtained neural activities with ultra-high density EEG, EOGs, and orbicularis oris EMGs under various speech conditions, including overt and covert speech. We then trained decoding models to predict spoken words and employed Explainable AI techniques to assess the neural contribution to EEG-based speech decoding. Finally, we examine the relationships between underlying neural signatures across speech conditions and discuss the versatility of EEG data from overt conditions.

## Results

### Real-time EEG decoding experiments

We conducted an experiment on the EEG-based classification of five words spoken under three speech conditions. These include overt speech condition (Fig. 1A *upper*. speech aloud), minimally overt speech condition (Fig. 1A *middle*. speech with minimal volume but without trembling of the vocal cords), and covert speech condition (Fig. 1A *lower*. chanting in the mind without vocalization). Participants in each speech condition freely manipulated a web interface (GMail) by indicating colors (green, orange, magenta, violet, and yellow) of web buttons to click. They were instructed to repeat the speech five times in one operation. We confirmed that the loudness of the speech decreased in the order of overt, minimally overt, and covert speech conditions (Fig. 1B). Using uhd-EEG, we measured participants’ EEG signals during speech. Fig. 1C shows the channels used for measurement, covering Broca’s area, auditory area, Wernicke’s area, and motor area. We selected this channel configuration to capture neural activities relevant to speech production. Additionally, we acquired EOG and EMG data alongside EEG measurements.

**Fig. 1.**
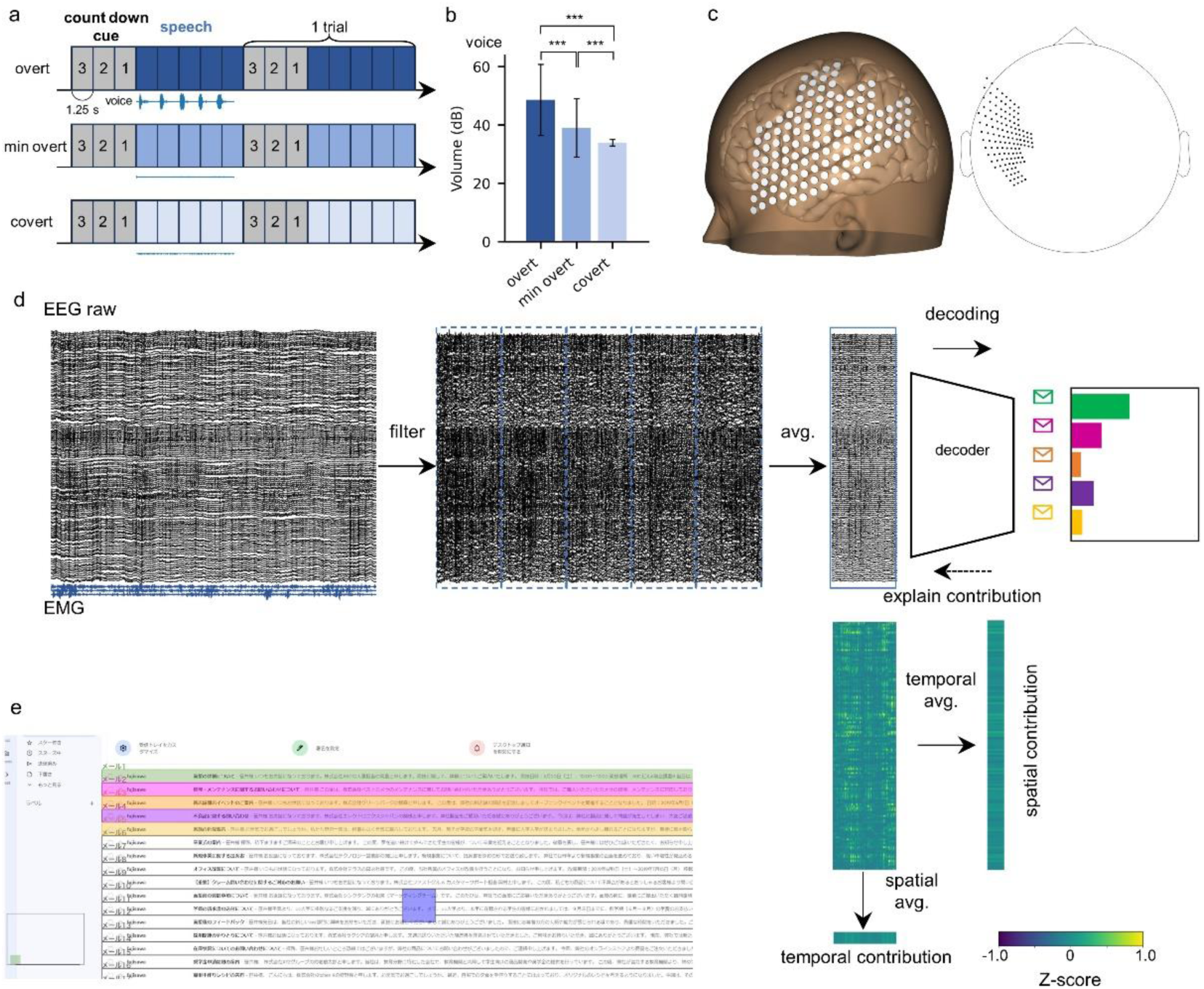
EEG-based decoding experiment. a. Experimental condition. Subjects were instructed to articulate the same word five times following a 3-countdown cue, with an interval of 1.25 seconds between each count and each speech. Speech production occurred in three modes: overt, minimally overt, and covert speech conditions. Overt speech entailed articulate pronunciation, minimally overt speech involved whispering quietly without extensive mouth movement, and covert speech required silent mental repetition of the words. b. Volume of each speech production. *** represents *P*<0.001 for Bonferroni corrected Wilcoxon rank-sum test after Kruskal-Wallis test c. EEG electrode layout. *left*: 3D, *right*: 2D d. Analysis pipeline. EEG data were acquired at a sampling rate of 256 Hz and were processed with a notch filter, common average reference, bandpass filter (2-118 Hz), and three myopotential removal adaptive filters. Subsequently, these data were segmented into 5 of 1.25-second intervals and averaged over the segments. The resulting EEG waveforms of 128 electrodes over 1.25 seconds were fed into the decoder to predict the spoken word among the five options. Spatio-temporal dynamics of EEG decoding contribution were obtained by tracing the gradients of inference in the decoder network and by calculating the integrated gradients. Spatial contribution was obtained as a temporal average of integrated gradients, while temporal contribution was determined by taking the inter-electrode average (*bottom right*). e. Online decoding experiment. We created an interface to control Gmail from EEG-based decoding. The interface uses five colors indicating web buttons to be clicked. By decoding a color spoken by a user, it can go back and forth on the Gmail web page, open an email of interest, generate candidates of replies, choose a reply, and send it. These candidate replies are created by ChatGPT based on the opened email and users’ past replies. The example image shows a participant controlling the interface by speaking a color corresponding to an email to be opened. Participants were instructed to produce each speech when the cue (*blue square*) appeared, and this was repeated 5 times at a fixed interval (1.25 seconds). The likelihood of the predicted word by the decoder is shown in the bar graph in the *lower left* corner, which can be seen by the user. The interface executed the command according to the decoded word (*i.e.*, the color with the highest likelihood).

We conducted decoding of the spoken words using collected data. First, we applied a notch filter to eliminate power line noise, a common average reference (CAR) for signal normalization, and a bandpass filter (2-118 Hz) to retain relevant frequency components. Additionally, we utilized an adaptive filter^29,30^ to remove predictable confounds from the EMGs at each time point. This was followed by averaging the signals over five consecutive speech segments of the same word to obtain denoised EEG waveforms (Fig. 1D *middle*). The denoised EEG signals were then fed into a word decoding algorithm, which was evaluated through 10-fold cross-validation using 100 instances of training and validation data (Fig. 1D *right*).

During online testing, we generated ensemble predictions by choosing four models with the lowest prediction errors on the validation data. We also calculated integrated gradients^31^ for successfully decoded trials to evaluate the contribution of EEG components to decoding. Spatio-temporal dynamics of the decoding contribution was obtained by taking temporal and spatial averages of the integrated gradients (Fig. 1D *bottom*).

### Adaptive filter suppresses EMG contamination

Muscle movements during recording likely contaminate obtained EEG signals. To evaluate these effects on EEG data, we measured three types of electromyograms (EMG): electrooculogram (EOG) and upper/lower orbicularis oris EMGs (EMG upper/lower). We confirmed that magnitudes (root mean square; RMS) for all EMGs in the overt condition were significantly higher than those in other conditions (Fig. 2A). While EMG lower in the minimally overt condition was significantly higher than that in the covert condition, other EMG channels showed no significant difference between these conditions. To quantify the information shared between EMG and EEG, we estimated mutual information for each pair of EMG and EEG channels.

**Fig. 2.**
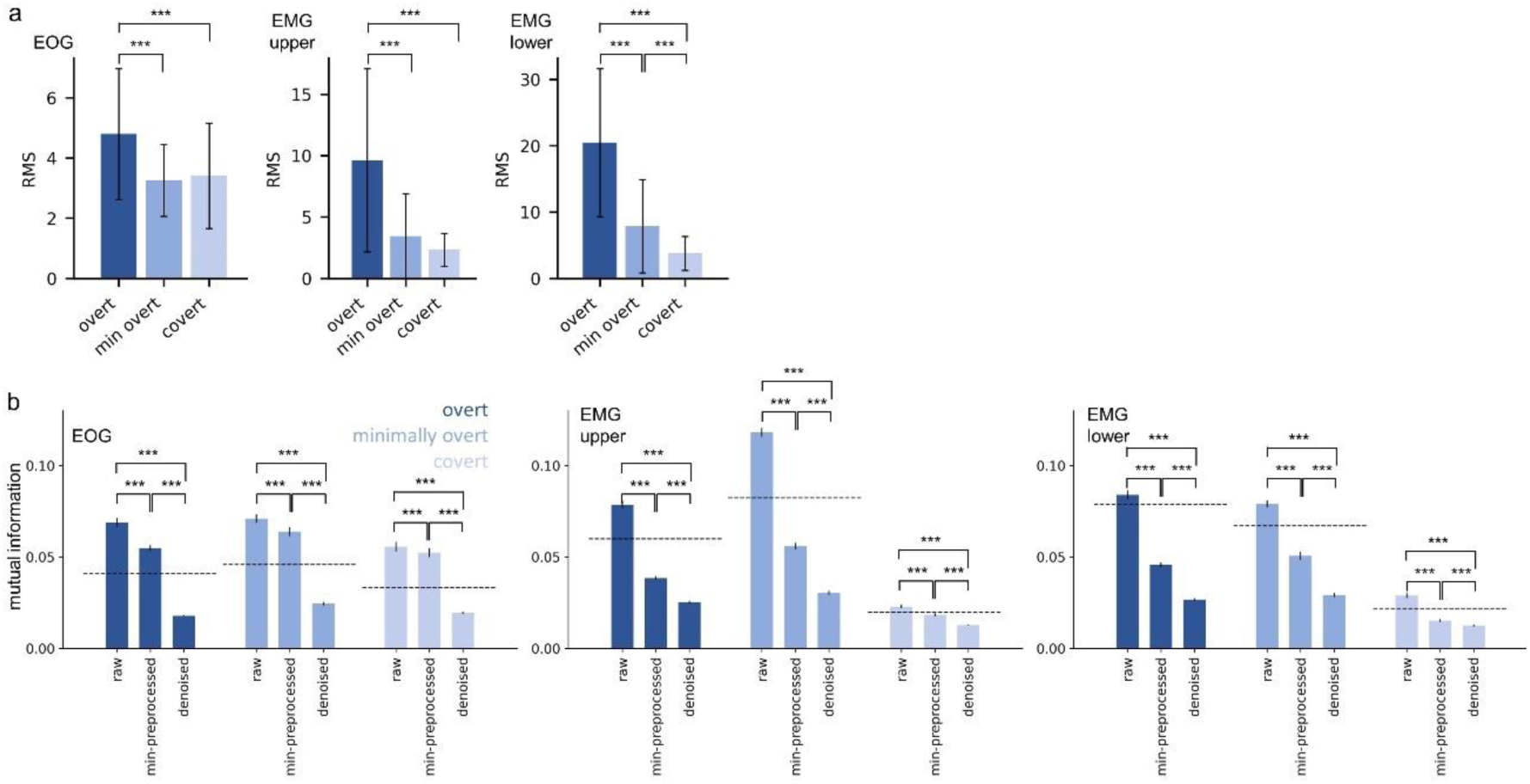
Artifact characterization. a. Root mean square (RMS) of EOG (*left)*, upper orbicularis oris EMG (*middle*), and lower orbicularis oris EMG (*right*) in overt, minimally overt, and covert speech productions, respectively. *** represents *P*<0.001 for Bonferroni corrected Wilcoxon rank-sum test, after Kruskal-Wallis test. b. Mutual information between EOG (*left*), upper orbicularis oris EMG (*middle*), and lower orbicularis oris EMG (*right*) at each stage of preprocessing. The mutual information between EEG and EMG during five consecutive speeches (1.25 sec × 5 = 6.25 sec, corresponds to single trial) was estimated using k-nearest neighbors methods and averaged across EEG electrodes. The bar graphs represent mean mutual information across trials and error bars represent SEM. Raw represents raw EEG data without preprocessing. Min-preprocessed is EEG after applying notch filter, common average reference (CAR), bandpass filter (2-118 Hz), and denoised is the residual EEG after subtracting linearly predicted components from EMG by adaptive filter. The mutual information was calculated separately for each speech production and averaged across electrodes. The dotted line represents the baseline mutual information when trial shuffling was performed within each recording session. *** represents *P*<0.001 for Bonferroni corrected Wilcoxon signed rank test, after Friedman test.

To verify the effectiveness of preprocessing in removing EMG from EEG signals, we compared the mutual information with EMG at each preprocessing step: raw EEG signals [raw], signals minimally preprocessed without the adaptive filter [minimally preprocessed], and fully denoised signals with the adaptive filter [denoised]. We confirmed that the mutual information decreases in the order of raw, minimally preprocessed, and denoised signals for all types of EMGs and speech production styles. Importantly, for EOG, the denoised EEG significantly decreased below the baseline (mutual information obtained by trial shuffling), and for upper and lower EMGs, the minimally preprocessed and denoised EMGs decreased below the baseline (see Fig. 2B), demonstrating that adaptive filter effectively suppressed EMG contaminations.

### Decoding model selection

To explore speech-related brain activity in EEG, we opted for a decoding model capable of predicting spoken words. Specifically, we compared EEGNet^32^, LSTM^33^, and SVM^34^, commonly used for EEG decoding. Although EEGNet and LSTM exhibited lower balanced accuracy for online tests (Table 2) compared to offline (validation) data (Table 1), they outperformed SVM in overt and minimally overt speech tasks. Furthermore, EEGNet showed remarkable online accuracy, with its peak performance achieved during minimally overt speech. Online decoding accuracies for covert speech were slightly higher than the chance for EEGNet and SVM, while it was lower than the chance for LSTM. Consequently, we chose EEGNet for subsequent analysis.

**Table 1.** Offline decoding performance.

|  | overt |  |  | min-overt |  |  | covert |  |  |
| --- | --- | --- | --- | --- | --- | --- | --- | --- | --- |
|  | EEGNet | LSTM | SVM | EEGNet | LSTM | SVM | EEGNet | LSTM | SVM |
| sub. 1 | **0.946** | 0.798 | 0.032 | **0.956** | 0.858 | 0.018 | **0.908** | 0.825 | 0.024 |
| sub. 2 | **0.975** | 0.925 | 0.697 | **0.954** | 0.812 | 0.195 | **0.927** | 0.830 | 0.148 |
| sub. 3 | **0.918** | 0.846 | 0.188 | **0.943** | 0.844 | 0.227 | **0.899** | 0.673 | 0.122 |
| avg. | **0.946** | 0.857 | 0.306 | **0.949** | 0.840 | 0.167 | **0.911** | 0.776 | 0.098 |

The balanced accuracy of EEGNet, LSTM, and SVM decoders for offline data in each speech condition, with each row showing the performance for an individual subject and the average for all subjects. The model with the highest accuracy for each speech condition is marked in bold.

**Table 2.** Online decoding performance.

|  | overt |  |  | min-overt |  |  | covert |  |  |
| --- | --- | --- | --- | --- | --- | --- | --- | --- | --- |
|  | EEGNet | LSTM | SVM | EEGNet | LSTM | SVM | EEGNet | LSTM | SVM |
| sub. 1 | **0.415** | 0.268 | 0.225 | **0.685** | 0.326 | 0.200 | 0.188 | **0.227** | 0.214 |
| sub. 2 | **0.712** | 0.407 | 0.479 | **0.564** | 0.306 | 0.200 | 0.184 | 0.104 | **0.200** |
| sub. 3 | 0.095 | 0.105 | **0.200** | **0.418** | 0.380 | 0.231 | **0.265** | 0.235 | 0.200 |
| avg. | **0.407** | 0.260 | 0.301 | **0.521** | 0.348 | 0.215 | **0.212** | 0.189 | 0.205 |

The balanced accuracy of EEGNet, LSTM, and SVM decoders for online data in each speech condition, with each row showing the performance for an individual subject and the average for all subjects. The model with the highest accuracy for each speech condition is marked in bold.

### Impact of electrode density on decoding

We assessed the impact of electrode density on decoding accuracy by using fewer electrodes than the original configuration. For this assessment, we utilized balanced accuracy of EEGNets trained with 4 distinct electrode subsets including 4, 8, 16, or 32 channels, respectively (Supplementary Fig. S1A). Offline decoding revealed that balanced accuracy significantly improved as the number of electrodes increased (Supplementary Fig. S1B, left). Similarly, for online decoding, balanced accuracy tended to improve with an increase in the number of electrodes, but significantly improved only during minimally overt speech tasks (Supplementary Fig. S1B, right).

### Distinct periods contribute to EEG-based and EMG-based decoding

We then investigated the relationship between EEG-based and EMG-based decoding for temporal periods on which EEGNet focused. To do so, we implemented a new EEGNet to decode words from minimally preprocessed EEG, and another from EOG, EMG upper, and EMG lower. Figure 3A shows representative time courses of integrated gradients. We obtained an average of integrated gradients over top-10 contributing EEG electrodes for all time points of data, while those of EOG, EMG upper, and EMG lower were shown as they are. Note that the integrated gradients were calculated by considering successfully decoded online trials. Figure 3B shows an example of the time courses of mean contributions for all trained EEGNets. Focusing on time points with higher electrode-average integrated gradients than their mean, we quantified the overlap rate of these featured periods between different data sets with the Jaccard index (Fig. 3C). As a result, we found a significant correlation between the upper and lower EMG signals for “magenta” and “violet”, and between minimally pre-processed EEG and upper EMG for “magenta” in minimally overt speech. A significant correlation between EOG and lower EMG was observed for “yellow” in overt speech. In contrast, no other significant correlations were detected, especially between denoised EEG and any EMG, demonstrating that EEGNets focus on distinct temporal periods when those signals are fed into them.

**Fig. 3.**
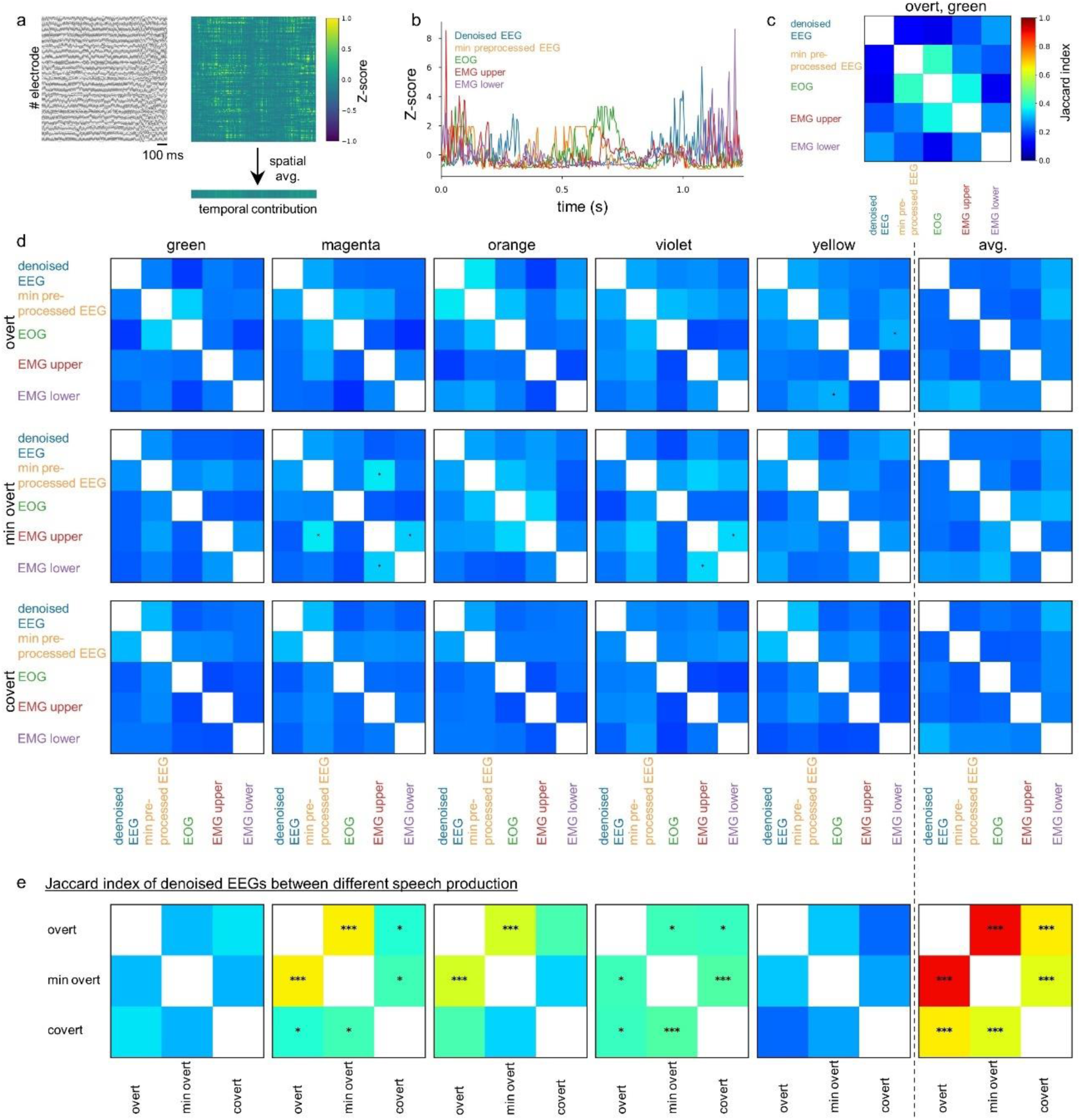
Distinct periods contribute to EEG-based and EMG-based decoding. a. Characterizing time periods contributing to decoding. (*left*) Denoised EEG for a representative trial. (*left*) Integrated gradient, the contribution of each EEG signal to word prediction, was calculated at each time point for each electrode. These values were then averaged across top-10 contributing electrodes to obtain temporal variations of contributions to decoding (*bottom right*). b. Exemplar temporal variations of contributions to decoding in a representative trial (“green” in the overt task): denoised EEG (*blue*), min-preprocessed EEG (*orange*), EOG (*green*), upper (*red*) / lower (*purple*) orbicularis oris EMG. c. Similarity (Jaccard index) matrix of the temporal changes in contributions of five distinct signals shown in B. d. Averaged similarity matrices across all trials are shown for each speech production and each word. Confidence intervals for the Jaccard index were estimated from the surrogate data.* represents *P*<0.05 for Bonferroni corrected bootstrap test. e. Similarity matrices between speech conditions. For denoised EEG signals, we calculated trial-averaged Jaccard indices of temporal variations in contributions to decoding between all conditions. * represents *P*<0.05 and *** represents *P*<0.001 for Bonferroni corrected bootstrap test.

Further analysis explored the congruence in featured periods across different speech conditions for EEGNets using denoised EEG for word prediction. We obtained Jaccard index matrices between the speech conditions and found significant correlations in featured periods for “magenta”, “orange”, and “violet” across overt, minimally overt, and covert speech conditions. We also observed an even stronger significant correlation for averaged featured periods over words between all speech conditions, highlighting that there are periods consistently featured in EEG-based speech decoding that are distinct from those in EMG-based decoding.

### Relationships of Distinct electrode-wide profiles between contributions to decoding and mutual information with EMG

We further investigate how EMG affects EEG-based decoding by comparing decoding contributions to EMG-EEG mutual information in an electrode-wise manner. Specifically, we quantified the electrode-wise contributions by taking temporal averages of integrated gradients (Fig. 4A) while obtaining electrode-wise EMG-EEG mutual information (Fig. 4C). Figure 4B shows mean contributions for each speech condition and produced word, as well as averaged contributions across all words. We found a high contribution of the temporal lobe across words, especially in overt and minimally overt speech, while spatial correlations between speech conditions showed no significant correlation (Fig.4D). We then examined the relationship between spatial distributions of contributions and those of EMG-EEG mutual information by calculating their spatial correlation coefficients (Fig. 4E). We found significant negative correlations between contribution maps in the overt condition and the mutual information maps. In contrast, no significant correlation was found for minimally overt and covert speech conditions.

**Fig. 4.**
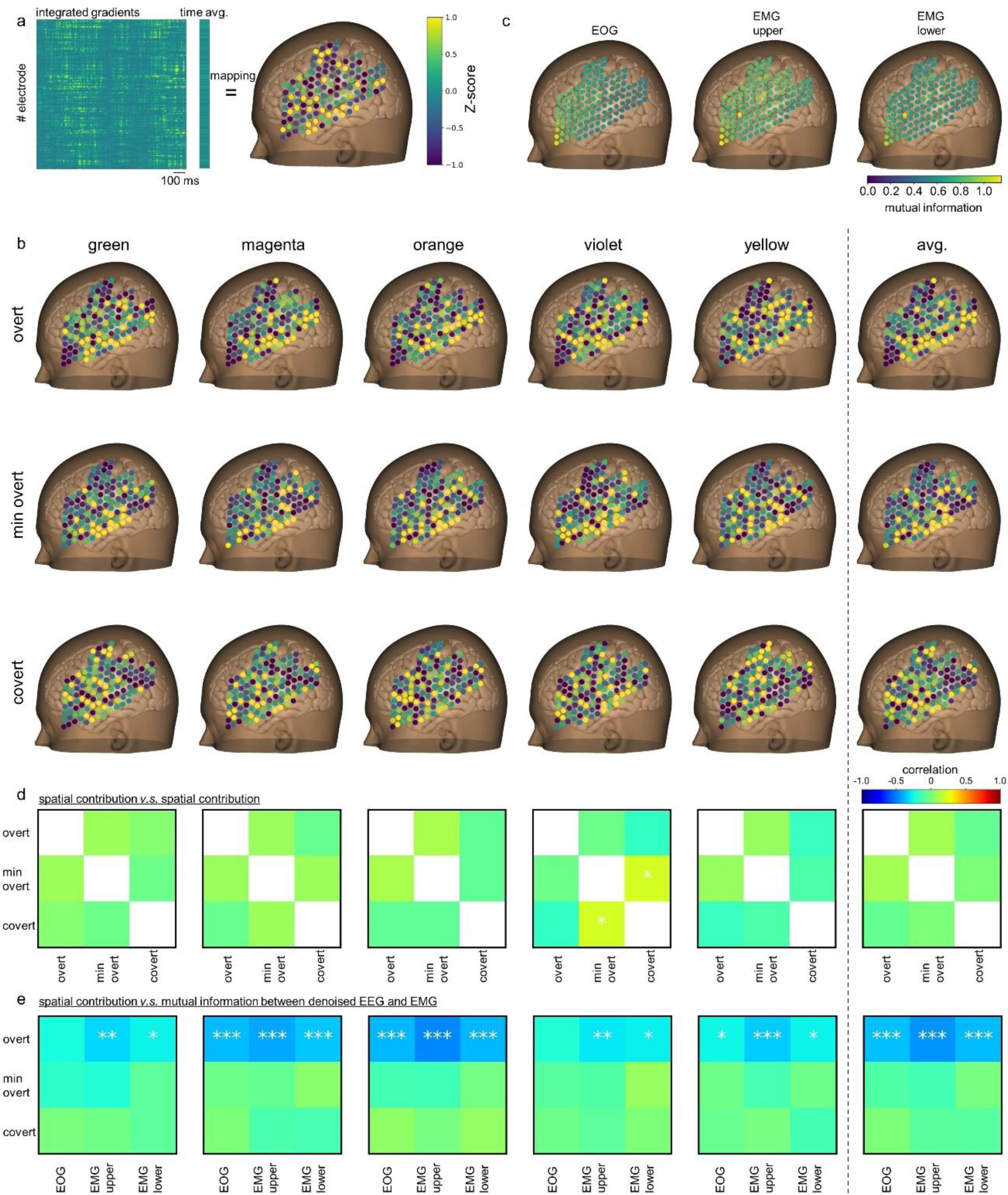
Electrode-wise contributions to decoding and mutual information with EMG. a. Evaluation of electrode-wise contributions to decoding. Integrated gradients, the contribution of decoding to word prediction, were temporally averaged and then standardized over electrodes to map onto the scalp surface. b. Electrode-wise contributions to decoding in each speech condition. These maps were obtained by averaging over trials for each word and all words, respectively. c. Electrode-wise mutual information with EMGs. Mutual information was estimated between a raw time series from an EEG electrode and that of EOG (*left*), upper orbicularis oris EMG (*middle*), or lower orbicularis oris EMG (*right*), respectively. d. Pearson correlation coefficients of spatial contribution between speech conditions for each word and the average of all words. e. Spatial correlation matrices between electrode-wise contributions to decoding and mutual information with EMG. * represents *P*<0.05, ** represents *P*<0.01, and *** represents *P*<0.001 for Bonferroni corrected Pearson’s correlation significance test.

### Adaptive filter enhances the contributions to decoding by speech-related brain regions

Adaptive filtering, suppressing the effect of EMG on decoding, may in turn lead to better contributions of neural activities to decoding, particularly for electrodes located within brain regions relevant to speech production. To investigate this hypothesis, we subtracted the electrode-wise Z-scored integrated gradients of the min-preprocessed EEG from those of the denoised EEG (Fig. 5A). Figure 5B shows changes in Z-scored integrated gradients for all speech conditions and produced words. We confirmed the suppression of contributions from a cluster of prefrontal electrodes showing high mutual information with EMG channels. Notably, we observed a consistent enhancement of contributions within auditory and language systems including Pars Triangularis and Pars Opercularis of the inferior frontal gyrus, precentral gyrus, angular gyrus, supramarginal gyrus, and superior temporal gyrus. Significant positive correlations were found for all pairs of conditions (Fig. 5C). We further investigated whether the adaptive filter artificially imposes spatial patterns of EMG contaminations, causing an overestimation of the correlations. To test this, we adjusted correlation coefficients by suppressing the contributions of electrodes according to their mutual information with EMGs (Supplementary Fig. S2). Significant correlations remain after the adjustment, demonstrating that EMG cannot explain the configuration of electrodes contributing to the correlation.

**Fig. 5.**
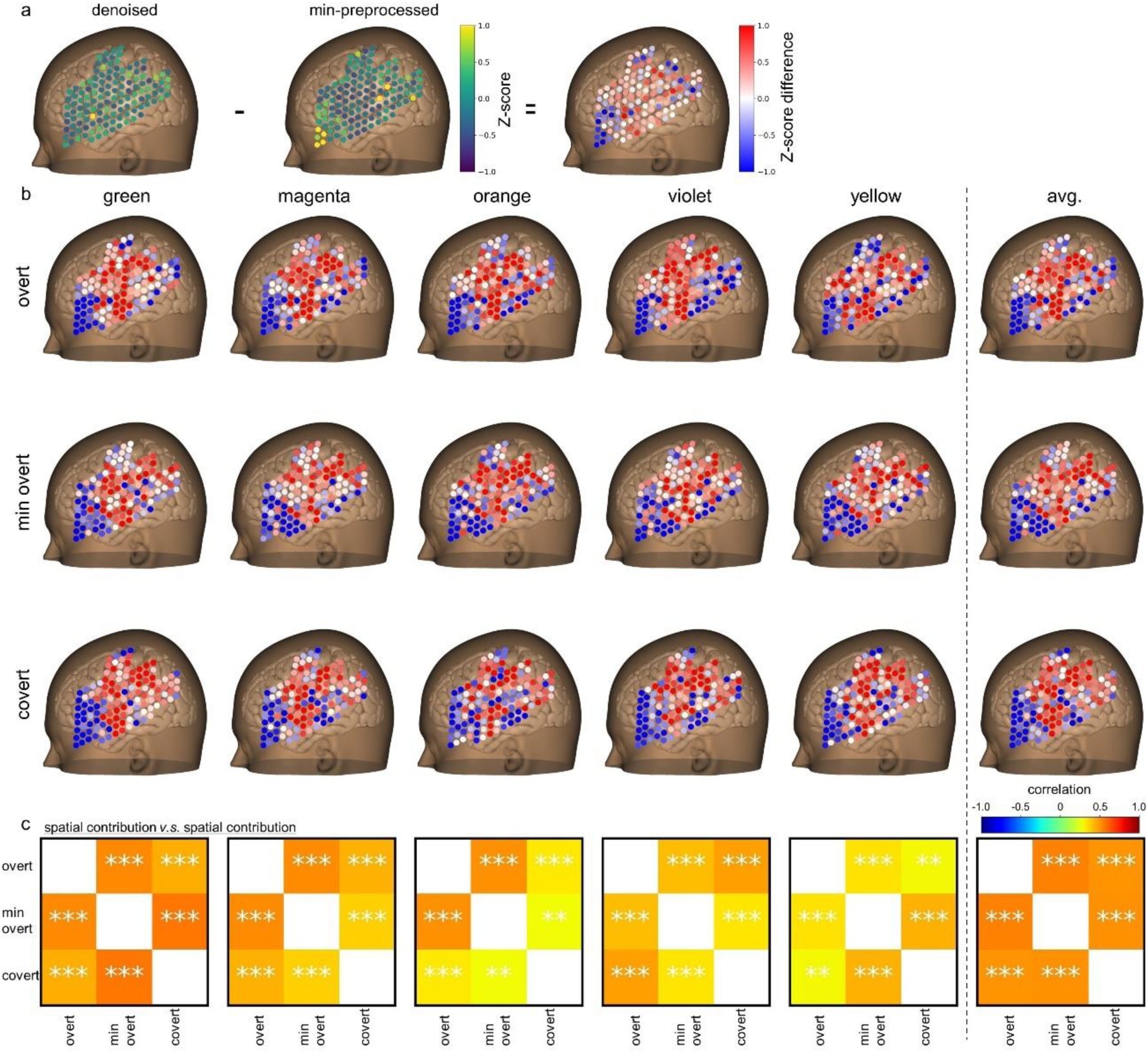
EMG removal enhances contributions of electrodes within speech-related brain regions to decoding. a. Contributions to decoding shifted by EMG removal through the adaptive filter. Differences in Z-scored contributions between min-preprocessed and denoised EEG data are mapped onto the scalp surface. b. Electrode-wise changes in contributions to decoding for each speech condition and mean changes over the conditions. c. Similarity of shifts in spatial patterns of contribution between conditions for each word and the average of all words. ** represents *P*<0.01 and *** represents *P*<0.001 for Bonferroni corrected Pearson’s correlation significance test.

## Discussion

We developed an online speech interface with EEGNet decoding overtly or covertly spoken words from neural activities recorded by uhd-EEG (Fig. 1). EEG data acquired during overt speech include facial muscular activities. By incorporating adaptive filters with a conventional preprocessing technique, we successfully suppressed EMG-EEG mutual information in the denoised data (Fig.2). Furthermore, we found contributions of distinct periods between EEG-based and EMG-based decoding (Fig. 3). Moreover, there was no similarity of electrode-wide patterns between contributions to decoding and mutual information with EMG (Fig. 4). Notably, we observed that EMG removal through the adaptive filter enhances the contributions to decoding by speech-related brain regions for all conditions (Fig. 5). These findings support the idea that combined with appropriate filtering processes, uhd-EEG is a useful device to decode spoken words based on underlying neural activities.

A silent speech interface, a device enabling speech communication without voices, would be an aid for those suffering from speech disabilities. Such interfaces may also improve the quality of daily-life communication in noisy environments or situations where vocal communication is not appropriate. Ideally speaking, covert speech, or imagined speech, is the least effort-intensive inputting method in using silent speech interfaces. While some studies using EEG have reported significant classification accuracy in offline decoding tasks for covertly spoken words^12,19–22^, investigations in online tasks^23^ are scarce. Indeed, we found that classification accuracy in the covert condition is high for offline speech, while it drops to around its chance level in online speech. On the other hand, minimally overt speech, where vocal sounds are maximally suppressed and hard to hear, showed the highest accuracy in both offline and online conditions. In addition, this condition effectively suppresses EMG contaminations compared to overt speech, which should improve neural contributions to decoding. Thus, we argue that minimally overt speech, rather than covert/imagined speech, is a practical solution for inputting EEG-based silent speech BMI.

EEG has been contributing to localizing relevant brain regions, especially with source localization techniques^35–39^. However, it gets harder and may become unreliable in the presence of strong confounds such as strong myoelectric potential generated by facial muscular movements. As such, most EEG-based speech studies have focused on neural events occurring prior to the onset of vocalization^40–43^. In contrast, we used uhd-EEG and localized electrode-wise contributions to decoding performance using signals denoised with adaptive filters. Notably, the enhancement in contributions via adaptive filter successfully outlined boundaries of featured electrodes along brain areas relevant to speech^40–46^ even without applying source localization. Furthermore, the boosted commitment of speech-related electrodes was consistently found between overt, minimally overt, and covert conditions. While previous studies have reported similarities in electrode-wise contributions between overt and covert conditions, their analyses lack artifact removal from muscle activity, which may ambiguate the neural source for decoding^19^. Uhd-EEG, in combination with appropriate denoising techniques, thus potentially facilitates EEG-based functional brain mapping.

The majority of EEG-based Brain-Computer Interface (BCI) investigations, including numerous studies focused on speech decoding, commonly employ an offline approach, where accuracy is assessed through post hoc analysis of EEG data acquisition^12,47^. This approach entails the risk of utilizing data with similar temporal structures and permits large model sizes that could be inefficient for online settings. Some studies also indicate that models optimized through offline processes may not align with optimal parameters during online usage due to factors such as the presence or absence of feedback^48^. Supporting this view, we also found a discrepancy between offline and online accuracy especially in the covert condition (Table 1,2), highlighting practical difficulties overlooked in studies focusing on accuracies in offline tasks. There are several potential causes of reduced online performance. Since brain states are continuously varying, distinct neural activities may appear despite speaking the same word, which could be even larger between separated sessions. For example, in this study there was an interval of 20-30 minutes between offline and online sessions due to the training of the decoder after offline sessions. Slight differences in task conditions between train and test sessions, such as the presence/absence of sensory feedback, may also amplify the variability of neural activities. Furthermore, reproducing speech while completely maintaining the vocal characteristics is difficult, especially in the covert condition where there is no sound feedback. Thus, robust implementation of decoders and interfaces absorbing these variabilities is crucial and should be tested in online settings to ensure practical applicability.

A promising strategy to develop BCIs is training neural networks through self-supervised learning (SSL)^15^. This learning method leverages unlabeled data by automatically generating labels from the data itself, enabling models to learn complex structures underlying the data without the need for external labels provided by humans. Given the variability of neural activities, such learning paradigms utilizing massive data would be effective in developing robust speech BCIs enabling naturalistic communication. However, since SSL needs paired data such as voices and concurrently obtained neural activities, it would be challenging to apply them as they are to EEG data collected during covert speech. Unlike covert speech tasks, overt speech tasks enable the objective retrieval of spoken words, as well as the coordination of muscles for speech production. As such, overt paradigms are practically appropriate for developing SSL-based BCIs, particularly in users’ natural speech settings. While the relationship of underlying neural dynamics between overt and covert speech conditions remains controversial, our findings indicate that uhd-EEG can measure consistent neural dynamics between these conditions. This view is further supported by previous studies, indicating shared neural mechanisms between overt and covert speech^19,21,49,50^. Future endeavors should focus on pre-training the model with extensive data from overt speech to enable transfer learning for covert speech data, which may become a more practical approach towards speech BCIs for real-world applications.

## Methods

### Participants

A total of 3 healthy adults (men; age range, 29–36 years) participated in this study. No participant had a history of neurological or psychiatric illness. Participants were initially given a briefing on the purpose of the study and experimental protocol, and if they approved informed consent, proceeded to the actual experiment. Our study was given ethical approval by the Shiba Palace Clinic Ethics Review Committee. The experiments were undertaken in compliance with national legislation and the Code of Ethical Principles for Medical Research Involving Human Subjects of the World Medical Association (the Declaration of Helsinki).

### Recording EEG and EMG

EEG was recorded from three healthy men (age range, 29-36 years). Before electrode placement, the hair was shaved with a shaver and then cleaned with alcohol tissue. The size of the head was measured with a measuring tape to determine the location of the CZ, and eight g.pangolin electrode sheets (16 electrodes per sheet, g.tec, Austria) coated with conductive gel (Elefix V, ZV-181E, NIHON KOHDEN, Japan) were attached at the marked positions. Electrode placement was determined around the language area of the left hemisphere (Fig. 1C). For the ground, the mastoid process behind the left ear was polished with Nuprep (Weaver and Company, US) and cleaned with alcohol tissue, and EMG/ECG/EKG electrode (Kendall^TM^, CardinalHealth, US) was placed. For EOG and EMG recording, the Kendall^TM^ electrodes were placed above and below the left eye and on two adjacent locations on the left upper orbicularis oris muscle and lower orbicularis muscle and were measured bipolarly.

### Preprocessing

EEG was acquired in real-time using the Python API of g.NEEDaccess (g.tec, Austria) at a sampling rate of 256 Hz. Notch filter (harmonics were also cut), common average reference, and bandpass filter (2-118 Hz) were applied. To reduce the EMG component contaminated in the EEG, the EEG signal linearly predicted from the EMG signals (EOG, EMG upper, and EMG lower) is subtracted from the EEG by an adaptive filter^29,30^. These filtered EEGs were then divided into 1.25-second segments and averaged over 5 speech segments. Thus, EEG waveforms of 128 electrodes for 1.25 seconds were obtained.

### Decoder architectures

Three decoding models were compared: CNN, RNN, and SVM.

CNN; EEGNet^32^ was used as the CNN model to decode spoken words from EEG. After temporal convolution with 120 ms kernel and spatial convolution, again temporal Separable Convolution^51^ was performed. Extracted spatio-temporal features of the EEG were used for classification with one fully connected layer.

RNN; It consists of 2 layers of bidirectional LSTM^33,52^ with 128 input and hidden units and one fully connected layer for classification of word labels.

SVM; Covariance between each electrode was calculated and projected to Tangent space to obtain features^34^. With these features as input, SVM (rbf kernel) was used to classify 5 words.

### Decoder training

At the beginning of the session, 100 trials of EEG data and speech labels were collected and used to train the decoding model. To deal with the possibility that subjects’ speech timing may shift back and forth during the online test, we sampled the EEG data with a time jitter of up to ±0.1 second from speech cue onset. To remove speech-independent noise between speech samples, a trial average of five consecutive speeches was taken to obtain a single training sample. For EEGNet and LSTM, we trained 100 epochs with a batch size of 16 samples and AdamW optimizer^53^ (weight decay=0.01) was used. The model weights of the training epoch with the lowest cross-entropy loss in the validation dataset were saved. For SVM, we iterated optimization until the change in the objective function was less than “tol”=0.001 (tolerance). 10 models were trained with 10-fold cross-validation.

### Ensemble decoding

During the online test, ensemble predictions were made using the top-k models (k=4), which had the lowest cross-entropy loss in the validation data set. For the ensemble prediction, the confidence of each predictor was Z-scored and averaged across predictors, and softmax was applied.

### Evaluation of decoding model

Because the number of trials among classes is not equal due to the data acquisition method, balanced accuracy was used to evaluate the performance of the decoding model (Table 1, 2).

### Brain-GMail interface

When the Brain-GMail interface was launched, the GMail home screen was shown. Five color-coded shades were overlaid on the top five emails. After a countdown of 3, 2, 1, a gray square appeared (turns blue after 5 speeches), blinking in 1.25-second cycles. Subjects freely chose one word from five different colors and spoke it timed to the appearance of these squares. After five speeches, the EEG data was sent to the decoding pipeline. The decoder’s confidence in each word was displayed as a bar graph. The email corresponding to the color with the highest confidence was opened. Next, the reply button was shaded “green” and the browser back button was shaded “magenta”. Again, after a countdown of three counts, the subject gave a speech of a word chosen from the five colors. If the interface opens an email that the subject has not indicated due to a decoder error, the user could return to the home screen by speaking “magenta”. If the interface correctly opens the email indicated by the subject, the subject spoke the word “green” to start the drafting of a reply to the email. Once the word “green” was decoded from the EEG, the content of the email was loaded into the ChatGPT 3.5 turbo^54^. ChatGPT was prompted to generate two positive and two negative replies each. The four reply patterns generated were displayed in “green”, “magenta”, “orange”, and “violet”, and the browser back button was assigned “yellow”. Subjects could indicate which reply they preferred by speaking the word, and if they did not like any of the replies, they could speak “yellow” to browse back. After one of the four colors of reply candidates was decoded, a sentence was selected according to the color. Next, the send button was assigned “magenta”, the reply button was assigned “green”, and the browser back button was assigned “orange”. Subjects could indicate the send button (“magenta”) if they had selected the intended sentence, or if they had made a mistake, they could start over again from sentence generation by indicating the reply button (“green”), or they could return to the home screen with the browser back button (“orange”). After the send button was selected, the reply button was assigned “green” and the browser back button was assigned “magenta”. When the Reply button (“green”) was selected, the ChatGPT began to generate the reply suggestions, and when the Browser Back button (“magenta”) was selected, the user was returned to the GMail home screen. In this way, an interface was developed in which the read, reply, and sending of GMail can be controlled by commands in five different colors specified with speech decoding.

### Speech task

The experiment was conducted with three speech styles: Overt (speech aloud), minimally overt (speech with minimal volume but without trembling of the vocal cords), and covert (chanting in the mind without vocalization). The subjects spoke synchronizing with cues that appeared every 1.25 seconds.

### Quantification of removal of EMG by mutual information

Mutual information for continuous variables was obtained using the method of estimating entropy from k-nearest neighbors distances^55^, which was obtained for each electrode for each EOG and upper/lower orbicularis oris EMG.

In Fig. 2B, the EEGs used to calculate the mutual information are compared with raw traces, traces minimally preprocessed with filters other than the adaptive filter (notch filter, CAR, and bandpass filter), and EEGs with all preprocessing including the adaptive filter. The baseline for the mutual information was determined by the mutual information when the combination of raw EEG and EMG was trial-shuffled within the recording session.

In Fig. 5C, the trial average of the mutual information for each raw EEG electrode was taken, and the amount of contamination of EMG into the EEG signals was quantified.

### Evaluation of decoding performance at different electrode densities

To predict the scaling of accuracy for electrode density, we hypothetically reduced the electrode density and compared decoding accuracy. Of the 128 total electrode channels, 32, 16, 8, and 4 channels were chosen and the decoding model was trained with the other electrodes masked, to obtain online balanced accuracy.

### Spatio-temporal structure of contribution to decoding

To visualize the spatio-temporal dynamics of the contribution to decoding (Fig. 3-5), integrated gradients^31^ were calculated according to the following formula

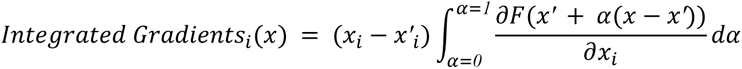

where the *i^th^* dimension of Integrated Gradients was computed. *x* ∈ *R^128×T^* denotes input EEG signals. T is the total number of time steps, which was 320 corresponding to 1.25 seconds at 256 Hz of Sampling rate. *x*′ ∈ *R^128×T^* is the EEG’s baseline, where each electrode’s average potential was used. *F* represents the EEGNet decoder, and *F*(*x*) is the inference confidence for the true label. 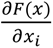 represents *i^th^* dimension of gradient of *F*(*x*).

### Statistical test

The significance level for all statistical hypothesis tests was set at *α = 0.05*. The Friedman test^56^, a nonparametric pairwise test or Kruskal-Wallis unpaired test, was used to test for differences between distributions of three or more groups. If the multiple comparison test showed a significant difference (*P < 0.05*, the Wilcoxon signed rank test, a nonparametric pairwise test or Wilcoxon rank-sum unpaired test, was performed. The P-value was corrected with Bonferroni’s method^57^ to control the false discovery rate when performing multiple comparisons. *** marked for corrected *P* value of *P*<0.001, ** for *P*<0.01, and * for *P*<0.05.

## Data availability

The raw data of EEGs / EMGs cannot be disclosed for ethical reasons related to the protection of personal information, but the codes used for the analysis are available in the following repository. https://github.com/arayabrain/uhd-gmail-public.git

## Acknowledgements

We thank Ryota Kanai for helpful discussions, Rousslan Fernand Julien Dossa for data collection, and Ippei Fujisawa for mock email generation. Special thanks to Anna Maria Hadjiev for her meticulous proofreading, significantly enhancing our manuscript’s quality. This work was supported by the JST, Moonshot R&D Grant Number JPMJMS2012.

## Author contributions

M.S.: Data curation, Formal analysis, Investigation, Methodology, Software, Validation, Visualization, Writing – original draft; Y.K.: Data curation, Resources, Investigation, Methodology, Writing – review \& editing; S.N.: Investigation, Methodology, Software, Writing – review \& editing; A.Y.: Investigation, Methodology, Software, Writing – review \& editing; M.I.: Investigation, Methodology, Software, Writing – review \& editing; M.S.: Data curation; K.T.: Data curation; S.S.: Conceptualization, Data curation, Funding acquisition, Project administration, Resources, Supervision, Writing – review \& editing.

---

**Supplementary Figure S1.**
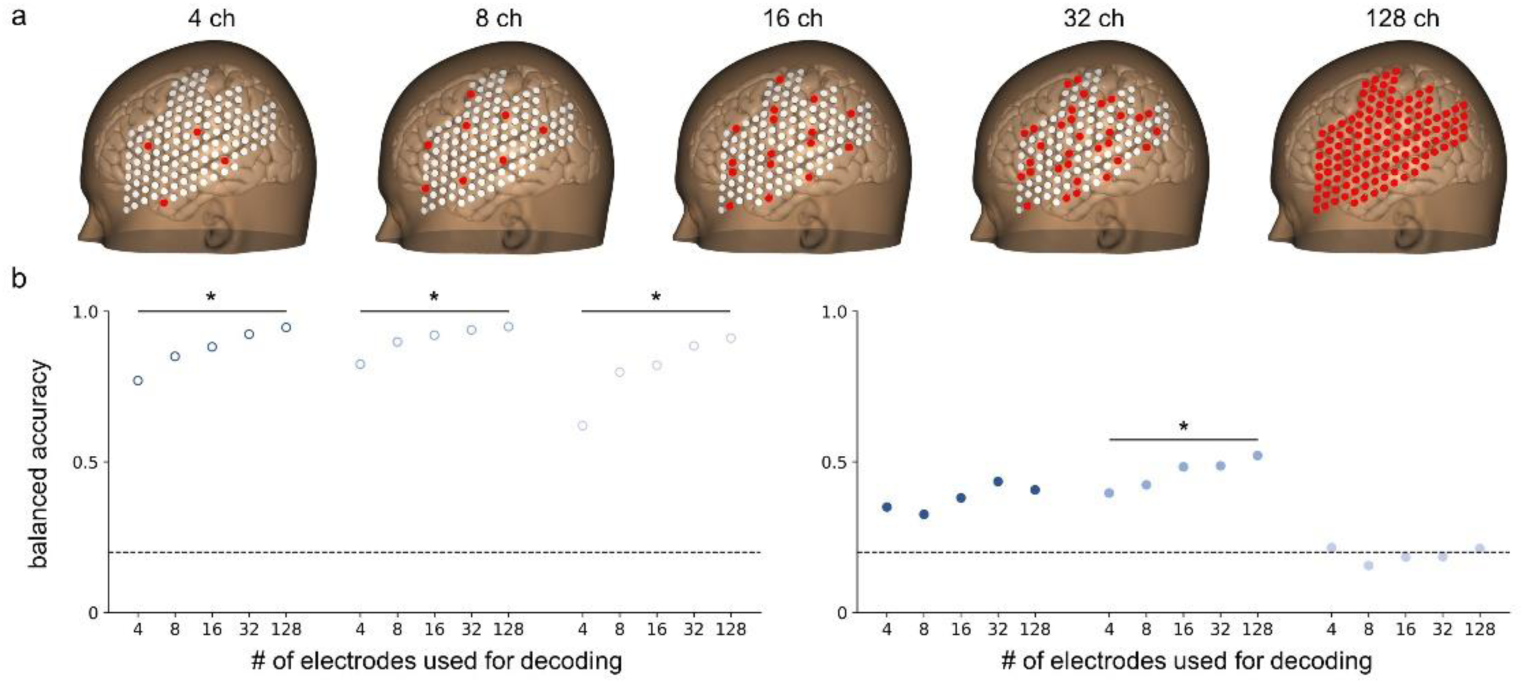
Relationship between electrode density and decoding accuracy. a. Electrodes used for decoding are shown in *red* while those not used (zero-filled) are shown in *white*. b. Balanced accuracies of EEGNet for offline (*left*) and online (*right*) trials were compared by hypothetically reducing the number of electrodes from 128 channels to 4, 8, 16, or 32. Conditions with a significant monotonically increasing trend of balanced accuracy for electrode density are marked with “*”. 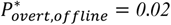, 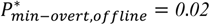, 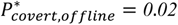, 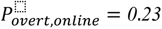, 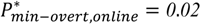, 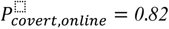, Kendall’s Tau test.

**Supplementary Figure S2.**
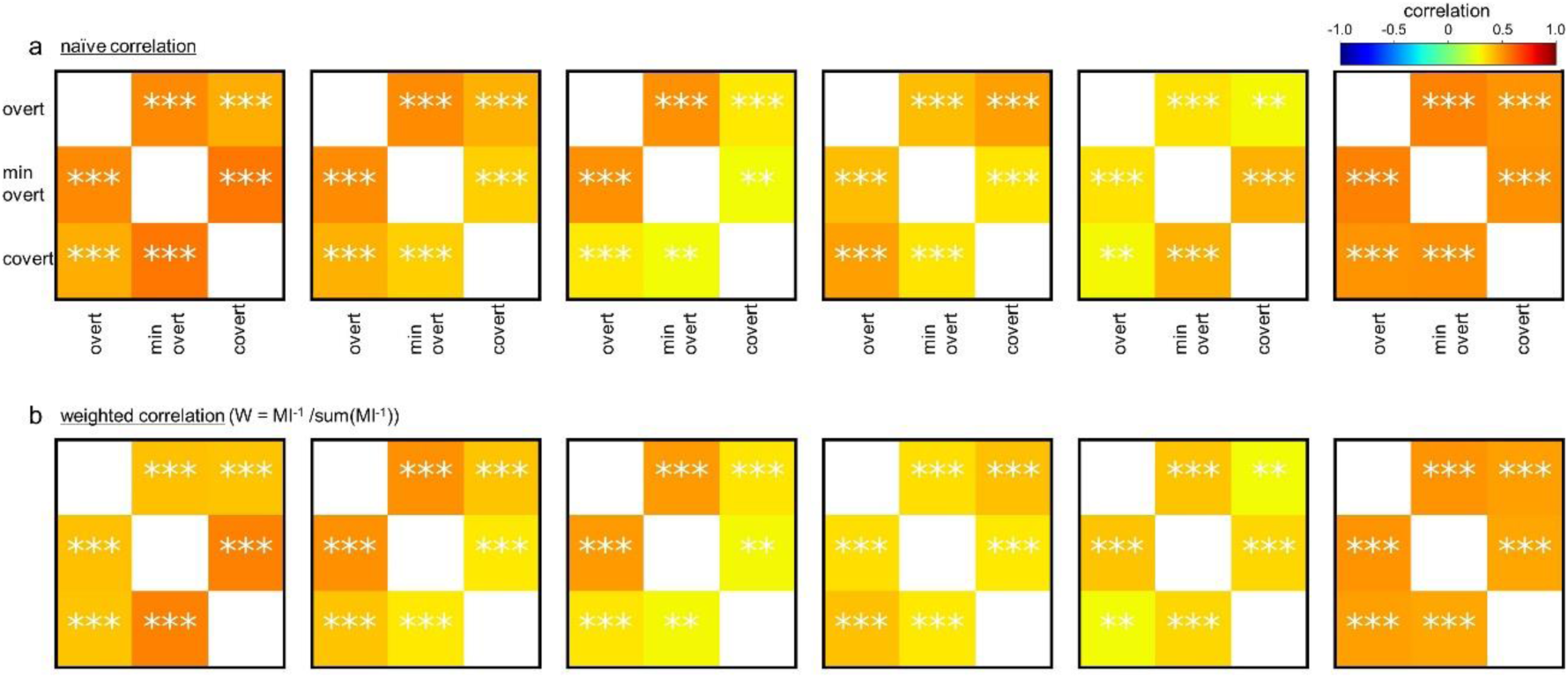
The correlation of shifts in spatial patterns of contribution between speech productions is not explained by the correlation of EMG patterns. a. Naive correlation of shifts in spatial patterns of contribution between conditions for each word and the average of all words. Same as Fig. 6C. ** represents *P*<0.01 and *** represents *P*<0.001 for Bonferroni corrected Pearson’s correlation significance test. b. Same as A, but for weighted correlation. The correlation coefficients were weighted by the reciprocal of the average of the mutual information between each EEG electrode and the EOG, EMG upper / lower in each speech production. ** represents *P*<0.01 and *** represents *P*<0.001 for Bonferroni corrected Pearson’s correlation significance test.

